# Defining the host dependencies and the transcriptional landscape of RSV infection and bystander activation

**DOI:** 10.1101/2025.03.26.645108

**Authors:** Sara Sunshine, Andreas Puschnik, Hanna Retallack, Matthew T. Laurie, Jamin Liu, Duo Peng, Kristeene Knopp, Matt S. Zinter, Chun Jimmie Ye, Joseph L. DeRisi

## Abstract

Respiratory syncytial virus (RSV) is a globally prevalent pathogen, causes severe disease in older adults, and is the leading cause of bronchiolitis and pneumonia in the United States for children during their first year of life [1]. Despite its prevalence worldwide, RSV-specific treatments remain unavailable for most infected patients. Here, we leveraged a combination of genome-wide CRISPR knockout screening and single-cell RNA sequencing to improve our understanding of the host determinants of RSV infection and the host response in both infected cells, and uninfected bystanders. These data reveal temporal transcriptional patterns that are markedly different between RSV infected and bystander activated cells. Our data show that expression of interferon-stimulated genes is primarily observed in bystander activated cells, while genes implicated in the unfolded protein response and cellular stress are upregulated specifically in RSV infected cells. Furthermore, genome-wide CRISPR screens identified multiple host factors important for viral infection, findings which we contextualize relative to 29 previously published screens across 17 additional viruses. These unique data complement and extend prior studies that investigate the proinflammatory response to RSV infection, and juxtaposed to other viral infections, provide a rich resource for further hypothesis testing.

**Importance:** Respiratory syncytial virus (RSV) is a leading cause of lower respiratory tract infection in infants and the elderly. Despite its substantial global health burden, RSV-targeted treatments remain unavailable for the majority of individuals. While vaccine development is underway, a detailed understanding of the host response to RSV infection and identification of required human host factors for RSV may provide insight into combatting this pathogen. Here, we utilized single-cell RNA sequencing and functional genomics to understand the host response in both RSV infected and bystander cells, identify what host factors mediate infection, and contextualize these findings relative to dozens of previously reported screens across 17 additional viruses.

## Introduction

Respiratory syncytial virus (RSV) is a ubiquitous respiratory virus that infects most children by two years of age, and reinfections are common [2]. RSV can lead to both upper and lower respiratory tract symptoms for all age groups, but remains a significant cause of global mortality for young children (< 5 years) and the elderly (> 70 years) [3]. While there are two FDA approved RSV vaccines for a subset of adults, and two monoclonal antibody treatments to protect infants from lower respiratory tract infection (LRTI), there remains no RSV-specific therapeutic for most infected patients. Notably, the two monoclonal antibodies (Palivizumab and Nirsevimab) that are available for prophylactic use are not clinically utilized for treatment of infection [4,5]. Due to the ongoing global health burden of RSV infection, it remains essential to continue investigating this important human pathogen.

RSV is a negative-sense, single-stranded RNA virus, and a member of the *Pneumoviridae* family [6]. RSV primarily infects respiratory epithelial cells and after viral entry, pathogen recognition receptors (RIG-I, MDA5) detect viral RNA and induce antiviral proinflammatory signaling cascades [7]. However, RSV encodes two nonstructural proteins (NS1 and NS2) that inhibit both interferon induction and signaling [8–10]. This dynamic interplay between the cellular antiviral response and RSV host-antagonism has been investigated in multiple transcriptional (bulk and single-cell), and proteomic studies *in vitro, ex vivo* and *in vivo* [8,11–15]. In addition to evaluation of the antiviral response to viral infections, perturbation based screens are commonly used to identify host factors required for viral infection. In the context of RSV infection, host dependency factors have been investigated using both RNAi [16] and haploid [17] screening.

To complement and extend these previous studies, we utilized both single-cell transcriptional profiling and genome-wide CRISPR screening to characterize the host response over the first 12 hours of infection and host dependencies for RSV infection. Our results identified host factors and pathways that are differentially expressed in infected and bystander activated cells, and revealed host dependency factors of RSV. While different viruses use a spectrum of infection and host subversion mechanisms, virologists have long sought to identify commonalities with the ultimate intention of discovering pan-viral therapeutic targets. Therefore, it is important to understand the degree to which RSV, or any other given viral pathogen, shares specific dependencies. Here, we compared and contextualized these results with respect to 29 publicly available contemporary CRISPR screens against other mammalian viruses (ie. SARS-CoV-2, HCoV-OC43, HCoV-NL64, ZIKV, EBOV) providing insight on shared and unique aspects of RSV host dependencies.

## Results

### Single-cell transcriptomics differentiates RSV-infected and uninfected bystander cells

To characterize the transcriptional response of RSV infected and uninfected bystander activated cells over the viral life cycle, we performed temporal single-cell RNA sequencing. A human lung epithelial carcinoma cell line (A549) was infected with RSV at an MOI of 0.3 (strain A2, consensus genome in File S1), and we performed droplet based single-cell RNA sequencing (10x Genomics) at four time points: 0 hrs, 4 hrs, 8 hrs and 12 hrs post infection (Fig. 1A). In parallel, heat-inactivated RSV was used to treat A549 cells for the same time points to control for differences between cells that underwent true infection compared to cells that came into extracellular contact with viral proteins and RNA. Furthermore, ambient nucleic acid is a potential confounder for droplet-based single-cell sequencing, especially for experiments with cytopathic viruses. These techniques may inadvertently capture cell-free nucleic acid, including viral RNAs or particles, therefore, it is essential to implement controls and a classification method to distinguish truly infected cells from those contaminated with ambient nucleic acid. Similar to our previous single-cell investigation of SARS-CoV-2 [18], uninfected cells (murine 3T3) were spiked-in following dissociation (Fig. 1A) to precisely assess ambient viral RNA quantities. Following quality control (methods), a total of 42,104 cells were captured across all time points, with a mean UMI count per cell of 18,676.

**Figure 1:**
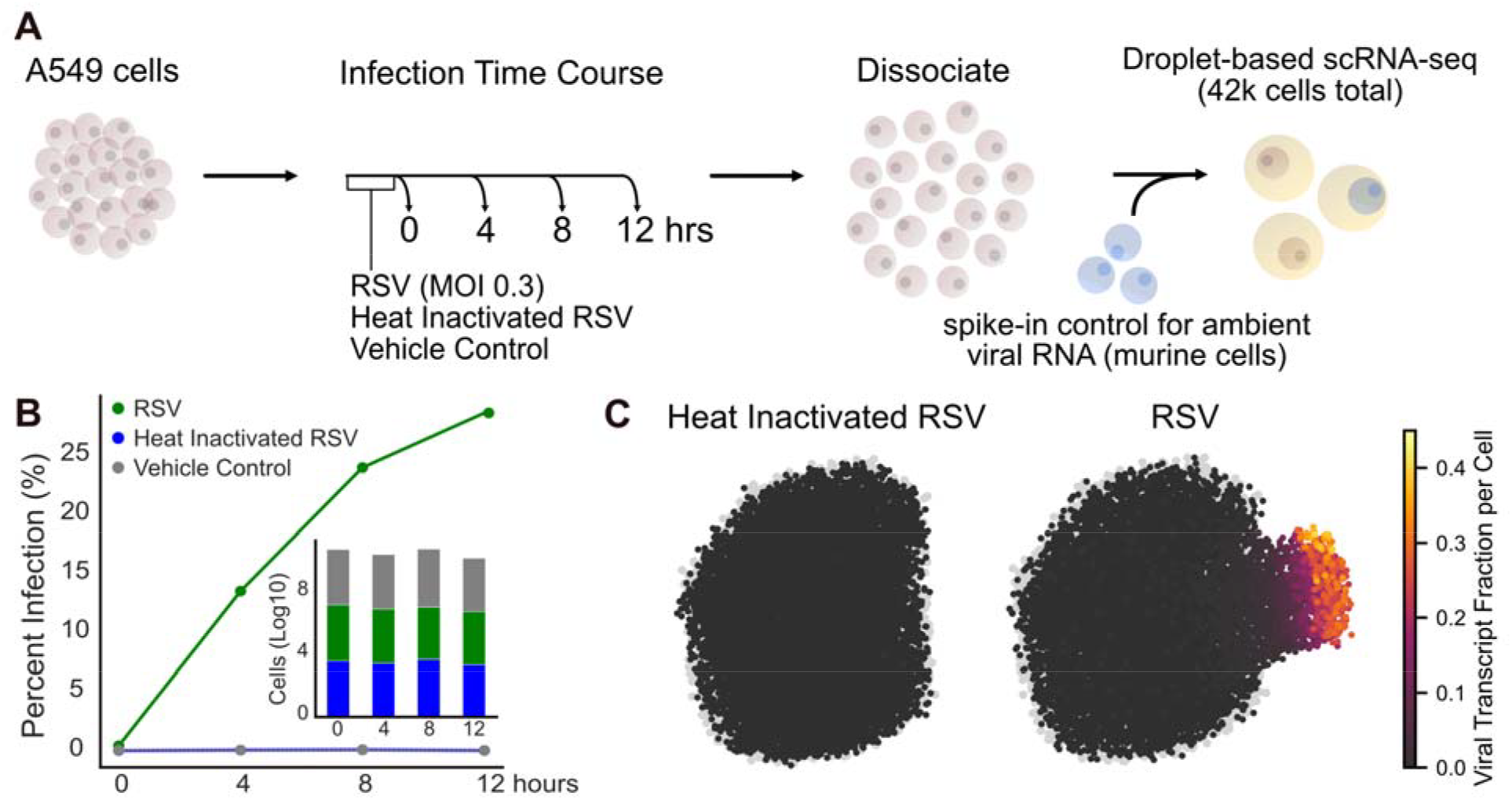
Experimental design and infection characterization. **A**. Experimental approach for characterizing the transcriptional response to RSV infection in single cells. **B**. We quantified the percent infection for each time point and condition. In the bar plot, the number of cells per time point and condition is displayed. **C**. Single-cell transcriptomes for all conditions were projected into UMAP space and cells treated with heat inactivated RSV or live RSV are colored by viral transcript fraction per cell and overlaid onto vehicle control treated cells (gray).

In order to investigate the transcriptional state of both infected and uninfected cells, we first had to identify infection status for each cell across all treatment groups and time points. We identified infected versus uninfected bystander cells by establishing a lower bound for ambient viral nucleic acid contamination in droplets from our spike-in cellular control (methods, Fig. S1A). Using this ambient viral RNA lower bound, the percent of infected cells was quantified for each condition and time point (Fig. 1B). Using the heat-inactivated RSV samples as an orthogonal control, the viral RNA lower bound was applied, and as expected, the vast majority (99.9%) of cells were classified as uninfected. With an MOI 0.3, 28.6% of cells were infected by 12 hours post inoculation. Additionally, we see a consistent number of cells for each time point and treatment group (Fig. 1B).

We next sought to understand the heterogeneity of viral transcript expression in single cells. On a per cell basis, while the viral RNA fraction of total cellular RNA per cell remained low for heat-inactivated RSV treated cells, as expected indicating no active replication in this group, we observed a heterogeneous viral RNA fraction for infected cells (Fig. 1C, Fig. S1B). Across all infected cells, we observe a mean viral RNA fraction of 8.2% and maximum viral RNA fraction of a striking 40.3%. The top 20% of infected cells accounted for 70% of the total viral RNA counts across all time points. While nearly a third of the cells were infected, these results suggest a small minority are producing disproportionately large quantities of viral RNA (Fig. 1C). Whether these same cells are disproportionately producing large numbers of viable virions remains an open question.

Over the course of infection and host cell antagonism, many viruses, such as SARS-CoV-2, employ host cell transcriptional shutoff. In contrast, cells with a higher fraction of RSV RNA had more total RNA counts (UMIs) than cells with less or no viral RNA (Fig. S1C). Upon removal of viral reads from the analysis, the remaining distribution of host UMIs were not substantially different from the uninfected/lowly infected population (Fig. S1D). Additionally, there was no correlation between viral fraction and host UMIs (Pearson’s r = 0.09). These results suggest that RSV lacks a broad host RNA shutoff detectable by single-cell RNA sequencing, unlike other viral pathogens (i.e. SARS-CoV-2), where host expression shut off is a hallmark of infection [18,19]. Whether RSV requires an intact host transcriptional context for viral production remains unknown. Regardless, the lack of host shutoff affords the opportunity to examine host transcriptional alterations within the infected population.

To investigate the transcriptional hallmarks of RSV infection, we focused on the time point featuring the largest number of infected cells, 12 hours, and projected cells from all three treatment groups into UMAP space (Fig. 2A-C). Analysis of the vehicle control and heat inactivated RSV treatment groups revealed a close overlap in UMAP space (Fig. 2A-B), suggesting the exposure of A549 cells to inactivated virus failed to provoke a measurable transcriptional response. Differential expression analysis (MAST [20]) confirmed that no genes were significantly differentially regulated (log2 fold change >2, padj <0.05) between the vehicle control and the heat inactivated control samples (Fig. S2A). For cells treated with live RSV, infected and uninfected bystander cells, as classified by the ambient viral RNA lower bound metric, were plotted separately in UMAP space (Fig. 2C). As expected, infected and uninfected cells occupied largely non-overlapping space in the UMAP projections. One of these clusters is dominated by truly infected cells, while the second cluster is composed of bystander cells (Fig. 2C).

**Figure 2:**
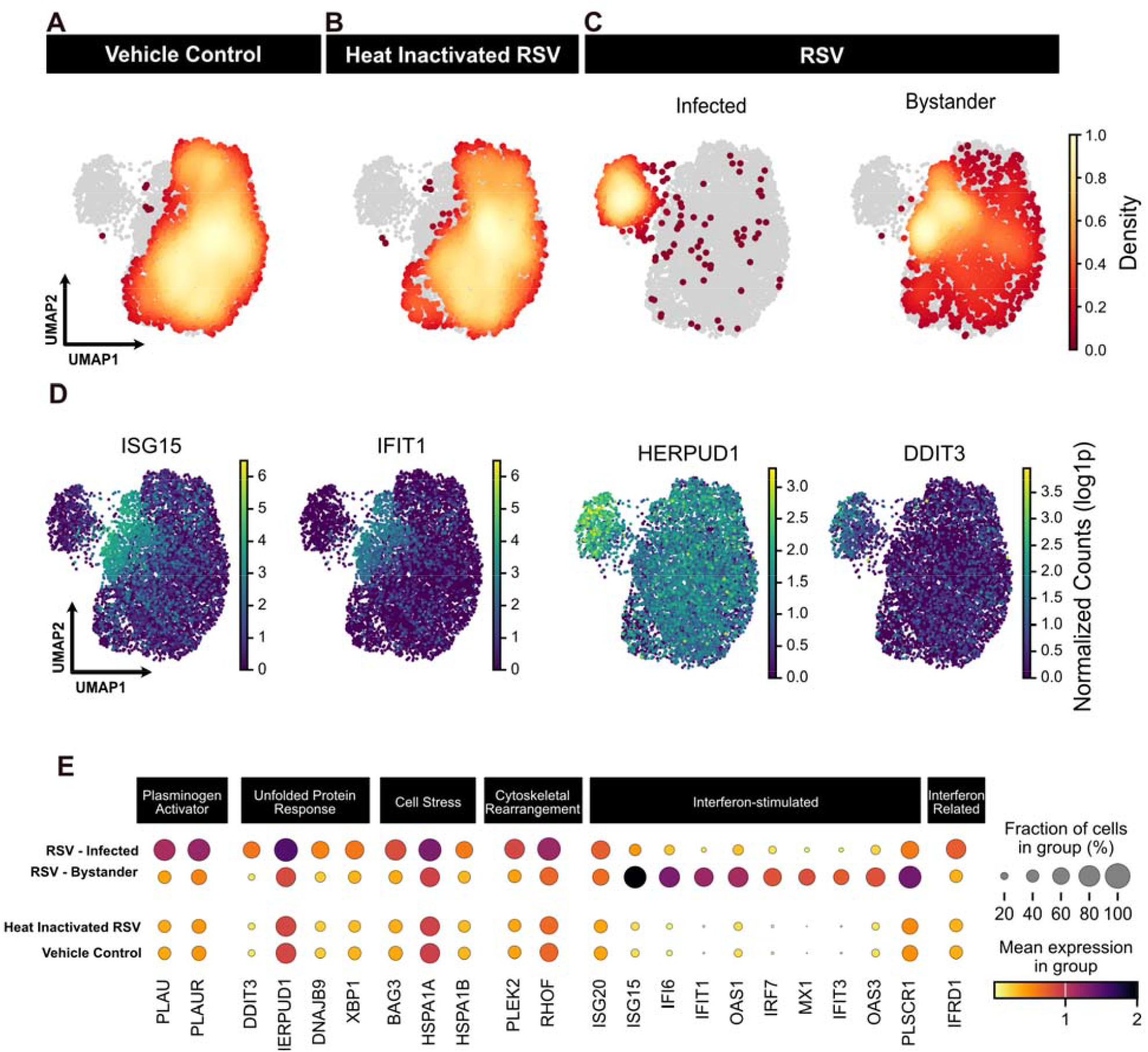
Single-cell transcriptome characterization 12 hours after RSV infection. **A-C**. Single-cell transcriptomes were projected into UMAP space and the density of cells for each condition were overlaid onto all cells (gray). The RSV infected condition (**C**) was further broken down by classification as infected or bystander cells. **D**. Transcriptional analysis of infected and bystander cells revealed distinct patterns in UMAP space, including differential expression for interferon-stimulated genes (ISG15, IFIT1) and genes associated with the unfolded protein response (HERPUD1, DDIT3). **E**. In the dot plot, mean expression for a subset of differentially expressed genes is plotted by condition. Dot size indicates fraction of cells in a group and color indicates mean gene expression.

To further characterize the transcriptional response in infected and bystander cells, we performed differential expression analysis (MAST [20]) between these cellular subsets and identified numerous genes spanning a variety of cellular processes that are differentially regulated (Fig. S2A, Table S1). Bystander cells featured a pronounced increase in abundance of canonical interferon-stimulated genes (ISGs), ISG15 and IFIT1 (Fig. 2D-E). Both ISG15 and IFIT1 are exemplars of well described interferon-stimulated genes with known antiviral functionality in the setting of SARS-CoV-2, HCMV and influenza infections [21]. Of note, ISG expression in bystander cells was detected as early as 4 hours after RSV treatment and differentially expressed 8 hours after infection (Fig. S2A-B). Conversely, ISGs were dramatically suppressed in the infected cells, consistent with virus mediated subversion of the host antiviral response (Fig. 2D-E).

RSV infected cells featured hallmarks of cellular dysregulation, including a statistically significant increase in genes associated with the unfolded protein response (DDIT3, HERPUD1, DNAJB9, XBP1), cellular stress (BAG3, HSPA1A, HSPA1B), and cytoskeletal changes (PLEK2, RHOF) (Fig. 2D-E). Additionally, plasminogen related (PLAU, PLAUR) factors, and the lysosomal cathepsin L protease (CTSL), were significantly upregulated in RSV infected cells (Fig. S2). Despite observing robust ISG downregulation in infected cells 12 hours post infection, ISG20 remains expressed in RSV infected cells. Notably, we observe in infected cells an upregulation of interferon-related developmental regulator 1 (IFRD1), a histone deacetylase-associated transcription factor, known to be upregulated in the setting of HPV infection that impairs proinflammatory activation [22]. It remains unclear if these expression patterns are due to higher baseline expression of ISG20 and IFRD1, or evasion of virally-induced ISG inhibition.

### Genome-wide CRISPR Screen Identifies Host Dependency Factors of RSV Infection

While single-cell differential expression efficiently identifies infection dependent transcriptional responses, such expression differences may not represent factors that are genetically required for RSV infection. To create a more comprehensive portrait of RSV infection and its genetic requirements, we performed two whole-genome CRISPR/Cas9 knockout screens (Fig. 3A). First, A549 cells were transduced with the GeCKOv2 library, and infected with RSV-EGFP for 24 hours. To enrich for host dependency factors, two different iterations of the screen were performed. For the first iteration (the “viability screen”), EGFP negative cells were sorted and cultured for 1 week, thus enriching for cells that were not infected at the time of the sort and remained viable after one week in culture. For the second iteration of the screen (the “early infection screen”), EGFP negative and EGFP-low cells were sorted 24 hours post infection and immediately harvested. For both screens, enriched sgRNAs, that presumably conferred protection from RSV infection, were PCR amplified and sequenced. Enrichment analysis was performed using the MAGeCK algorithm [23] to rank genes and calculate the MAGeCK enrichment score for each screen (Table S2).

**Figure 3:**
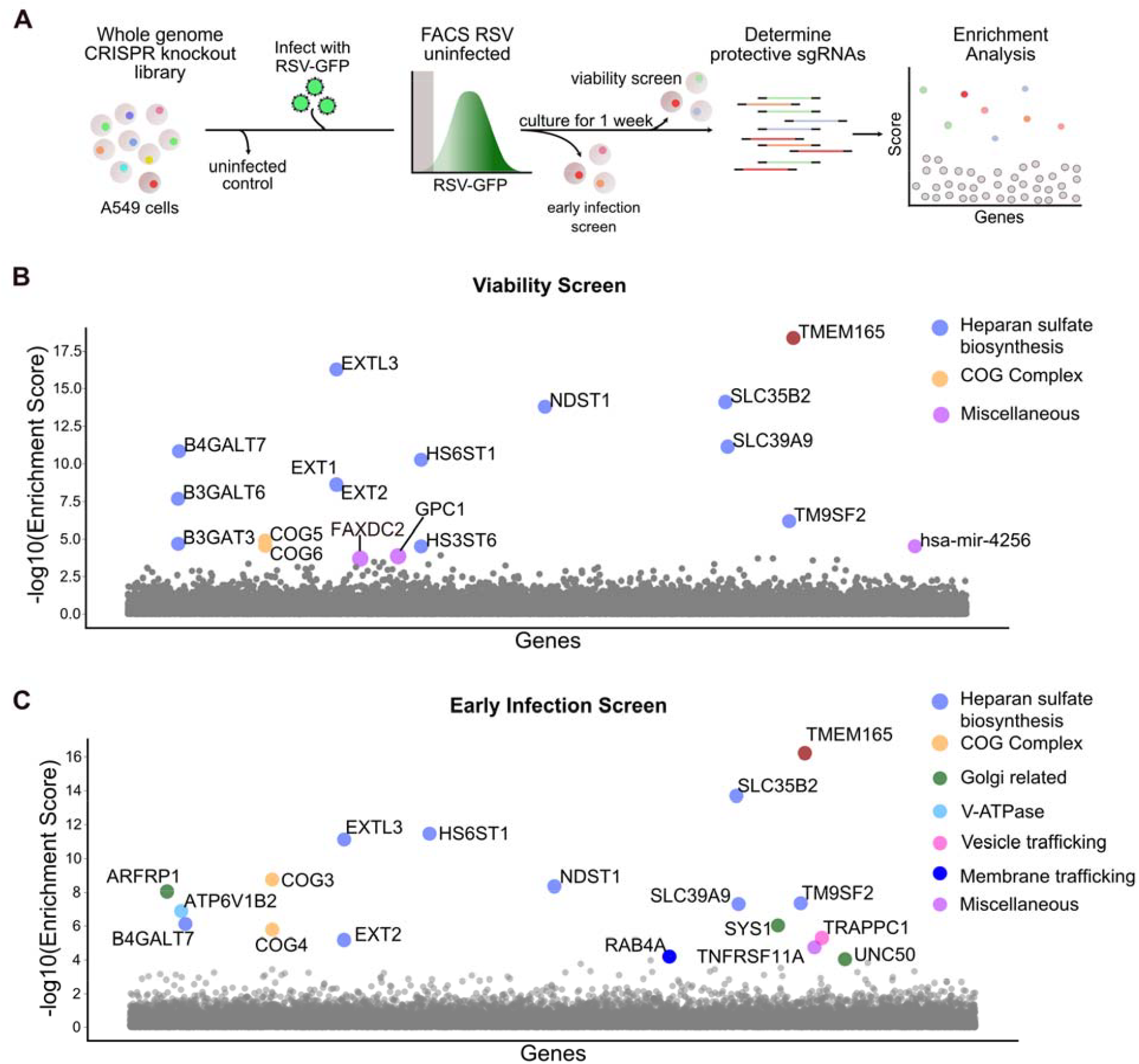
Genome-wide CRISPR/Cas9 knockout screen in human A549 cells for identification of host factors important for RSV infection. **A**. Experimental design for CRISPR/Cas9 screens. **B-C**. Gene enrichment scores for the “Viability Screen” (**B**) and the “Early Infection Screen” (**C**). Enrichment scores were determined using the MAGeCK algorithm [23] and gene hits are colored by function. The top hit for both screens, TMEM165, is highlighted in brown. A complete list of enrichment scores and genes can be found in Table S2.

First, the viability screen identified genes involved in heparan sulfate biosynthesis, the conserved oligomeric Golgi (COG) complex, and retrograde transport at the trans-golgi network (Fig. 3B). Notably, the top hit was TMEM165 (Transmembrane Protein 165). This protein is localized to the Golgi apparatus (Human Protein Atlas [24]), and is functionally thought to play a role in protein glycosylation. Knockout of TMEM165 has also been shown to reduce surface expression of heparan sulfate, and heparan sulfate proteoglycans are well known as viral attachment factors [25,26].

In concordance with the viability screen, TMEM165 was the most enriched gene for the early infection screen. For the early infection screen, along with heparan sulfate and COG complex related genes, additional host factors impacting RSV infection were identified, including those related to the Golgi (ARFRP1, SYS1, UNC50), V-ATPase (ATP6V1B2), vesicle trafficking (TRAPPC1) and membrane trafficking (RAB4A) (Fig. 3C). A comparison of both screens revealed substantial concordance in the top ranked genes (Fig. S3A). Differences between the two screens likely reflect the separate timescales (one week vs. 24 hrs). Furthermore, the viability screen may select for genetic perturbations that both prevent infection and prolong cell life/proliferation, while the early infection screen may be more specific to perturbations that prevent immediate infection, as opposed to prolonged survival. Alternatively, the early infection screen may be less stringent than the viability screen, possibly enabling the identification of host factors with partial phenotypes that may not survive in the viability screen.

To further identify pathways associated with our list of enriched genes, we subsequently performed gene ontology, pathway and process enrichment analysis using Metascape [27]. Significantly enriched pathways include heparan sulfate and glycosaminoglycan biosynthesis, cytosolic transport, retrograde transport at the transit-golgi-network and endoplasmic reticulum to Golgi vesicle-mediated transport (Fig. S3B). Notably, these pathways have been implicated as entry requirements for other viruses [28–30]. These findings confirm and extend multiple reports of heparan sulfate biosynthesis and trafficking associated factors as important for RSV infection, despite different screening methods and cellular models [16,17]. Comparing the single-cell differential expression results with the CRISPR knockout screen revealed no overlap, highlighting the importance of pursuing orthogonal approaches to elucidate the impacts and dependencies of viral infection.

CRISPR knockout screens for viral dependency factors have become increasingly useful in virology [31], however few studies have contextualized screening results across disparate viral species. Despite the difficulties in aggregating different studies, understanding both the commonalities and the differences between viruses and their reliance on specific factors may help capture the range of pathways and targets they subvert. This knowledge may also assist with future pursuits for pan-viral therapeutics by identifying overlapping factors. To investigate similarities and differences between RSV and other viruses, we curated a collection of 29 published genome-wide CRISPR knockout screens for 9 different viral families (*Arenaviridae, Coronaviridae, Filoviridae, Flaviviridae, Herpesviridae, Orthomyxoviridae, Picornaviridae, Poxviridae* and *Reoviridae*) (Table 1). To compare results across these different screens, we downloaded and processed datasets using a single statistical framework (methods). It is important to note that these published reports utilize a variety of cell lines, screening selections and sgRNA libraries, which may confound interpretation.

**Table 1:**
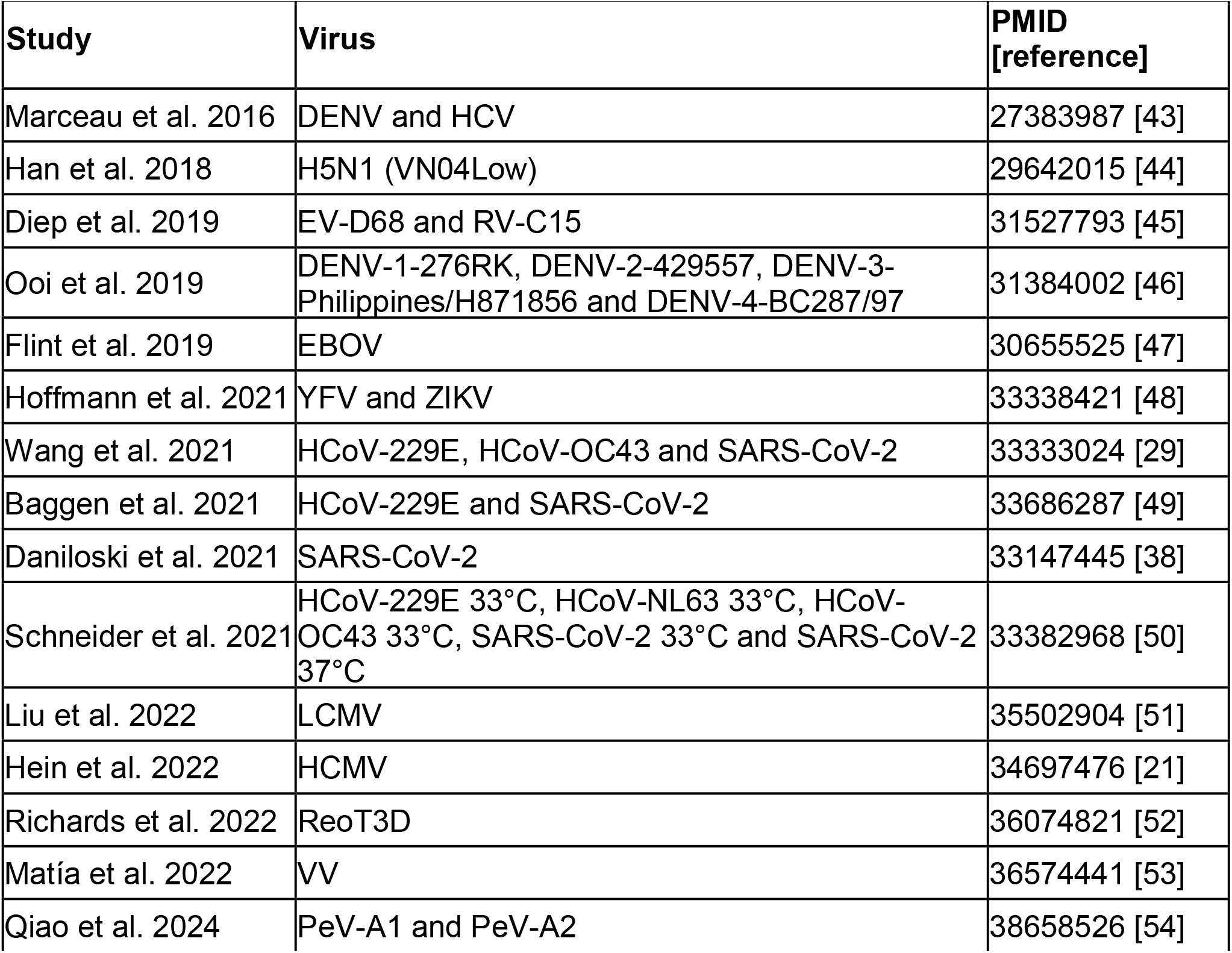
Published CRISPR screens for viral dependency factors utilized for comparative analysis.

Host factors were selected that were top hits in at least one or more screens (MAGeCK analysis method, FDR < 0.01). With the resulting 279 genes, hierarchical clustering was performed by gene and by CRISPR screen, and visualized by heatmap (Fig. 4, fully labeled heatmap in Fig. S4, Table S3). In several cases, viruses from the same family yielded substantial overlap. In particular, numerous members of the oligosaccharyltransferase (OST) complex and ER membrane protein complex (EMC) were a common hit amongst flavivirus screens, including DENV, YFV, and ZIKV (Fig. 4). The EMC complex plays a role in co-translational insertion of transmembrane proteins in the ER, which is a well appreciated facet of the flavivirus life cycle [32–35]. Likewise, several screens against coronaviruses yielded shared hits mapping to the conserved oligomeric Golgi (COG) complex, which serves multiple functions in membrane trafficking and glycosylation [36,37]. Notably, several members of this complex were also shared with our RSV screens. While many examples of sharing amongst subsets of viruses were observed, this analysis also makes clear that there exist no obvious gene hits that were “pan-viral.” Many viruses revealed distinct clusters of hits that were only seldomly shared, some of which may have depended on cell type. For example, the screen against LCMV [28], and a single SARS-CoV-2 screen [38], both in A549 cells, clearly surfaced multiple components of the vacuolar(H+) ATPase, which is responsible for endosomal acidification and viral entry. These aggregated results, spanning 29 screens and 17 viruses, suggest evidence for a multitude of virus specific and cell type specific infection and replication determinants, with only limited sharing amongst small subsets of often related viruses.

**Figure 4:**
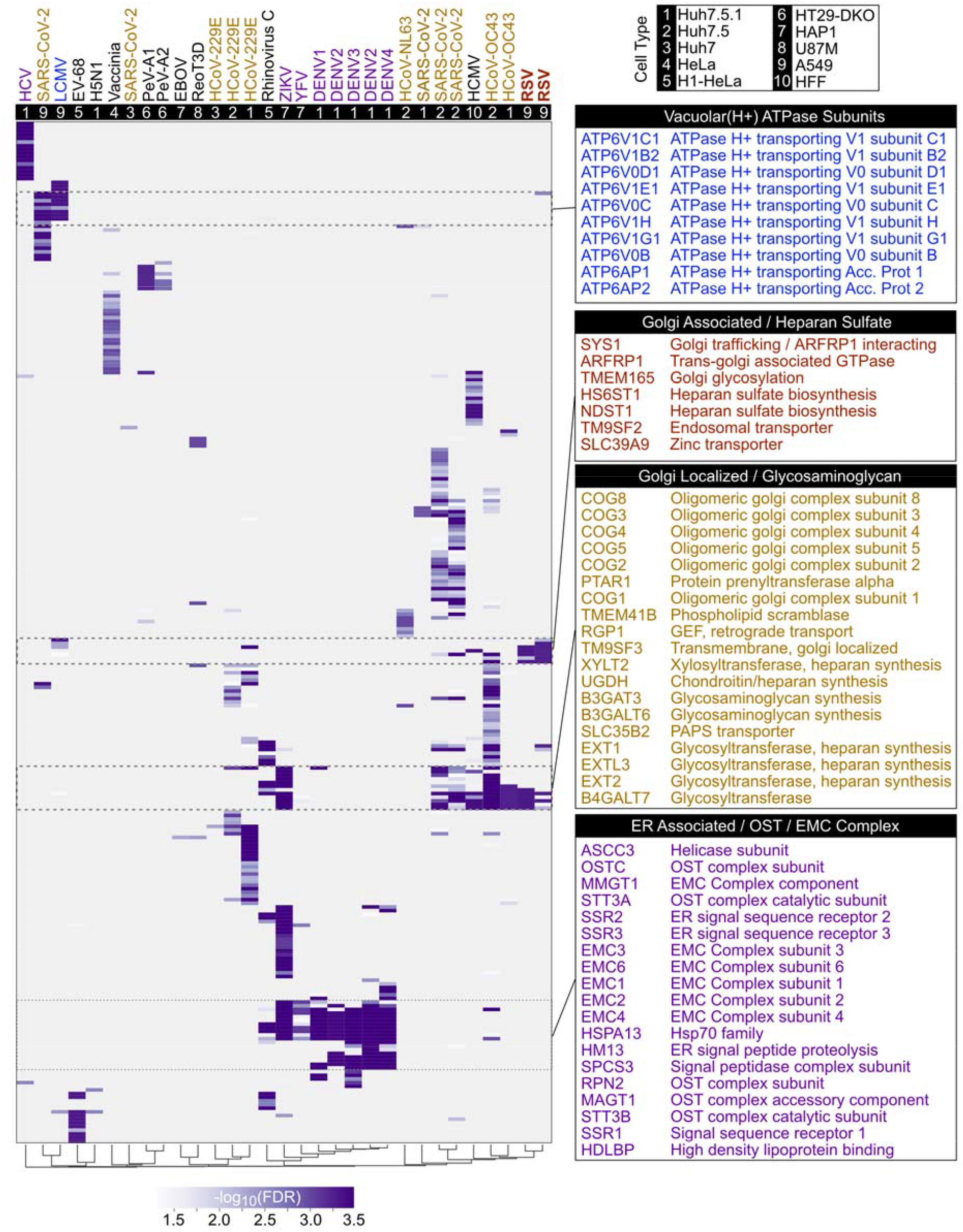
Comparison of RSV screening results with 29 published CRISPR/Cas9 screens for viral dependency factors. Host factor genes that were identified in at least one CRISPR screen using the MAGeCK analysis method (FDR < 0.01) were visualized on a heatmap for 29 published screens in addition to the two CRISPR screens from this study. Rows represent genes and columns represent CRISPR screen datasets. For genes that passed a threshold of -log10(FDR) > 1.3, heatmap blocks are colored by the -log10(FDR).

## Discussion

Although recent advances in vaccine technology hold great promise, RSV remains a public health burden without RSV-directed treatments available for infected populations. Therefore, it remains essential that we understand the host response and cellular factors required for infection. In this study, we utilized two unbiased methods to better characterize the underlying biology of RSV infection: single-cell RNA sequencing and CRISPR/Cas9 screening. These two techniques provided complementary, non-overlapping insights into the cellular biology of RSV.

It is well recognized that viral infection is a dynamic, heterogeneous process where both infected and bystander cells may contribute to the antiviral response [18,21]. The timing and magnitude of this antiviral response are crucial for viral clearance, and the first cells to get infected set the stage for this. To characterize the transcriptional response and viral antagonism during this critical period, we performed temporal single-cell transcriptional profiling every four hours for the first twelve hours of infection. Our transcriptional profiling reveals a dynamic process with intermediate transcriptional states, in addition to impacts on bystander cells. The analysis of our single-cell transcriptional data was significantly aided by rigorously controlling for ambient, cell-free RNA, which is a known confounder of droplet-based single-cell sequencing experiments. Using a spike-in control cell population allowed for accurate differentiation and comparison of infected and uninfected cells.

We observed that infected cells with the most viral RNA have more total UMIs per cell than uninfected cells, but do not dramatically differ in the total human UMIs per cell. While it is well understood that RSV host antagonism occurs through production of viral nonstructural proteins NS1 and NS2, these findings show that at the single-cell level, antagonism does not induce global host transcriptional shutoff. This result is similar to what is observed during influenza infection [39], and is in stark contrast to other RNA virus infections (i.e. SARS-CoV-2) [18]. This suggests that RSV possesses unappreciated host transcriptional dependencies.

Our transcriptional analyses revealed that the innate immune proinflammatory signaling cascades commonly associated with RSV infection are primarily observed in uninfected bystander activated cells. RSV infected cells showed strong downregulation of most ISGs, presumably through modulation by virally-expressed antagonists (NS1 and NS2). Consistent with a concerted effort to shut down the host antiviral response, we observe upregulation of Interferon Related Developmental Regulator 1 (IFRD1), a gene known to deregulate the NFκB pathway and lead to a decrease in proinflammatory signaling. Prior studies have shown that HPV upregulates IFRD1, suggesting this is a broad viral mechanism to impede the immune response [22]. Notably, one antiviral gene with known RNase properties, ISG20, remained at the same level or perhaps slightly more than the expression observed in bystander cells. ISG20 is in contrast to other classic ISGs, such as ISG15, which was significantly diminished in infected cells. Expression of ISG20 is known to inhibit replication for many other RNA viruses (i.e. IAV, VSV) [40,41]. In the context of RSV infection, where ISG20 expression appears undiminished, it remains unclear whether there exists uncharacterized host dependencies or if RSV is unperturbed by its action.

Resolution at the single-cell level further enabled us to identify modulated pathways specific to RSV infected cells, and our spike-in controls allowed us to distinguish these from bystanders. We observed upregulation of genes associated with the unfolded protein response, cell stress and plasminogen receptor activation. While prior bulk transcriptional studies have shown upregulation of these pathways [11], here, we show this signal is specifically expressed in the infected cellular population.

Single-cell transcriptomics provided unique information about the response in infected and bystander cells during RSV infection, however, required host factors may elude detection for a variety of reasons. For example, transcriptional levels of host factors may not change over the course of infection, or expression may be too weak or transient to measure reliably with this approach. As a complementary method to investigate the host determinants of infection, we performed two whole-genome CRISPR/Cas9 screens. These screens differed experimentally on timing and processing, and produced highly concordant results. For both screens, the most highly enriched host factor that provided protection from infection was Transmembrane Protein 165 (TMEM165), a protein known to be involved in trafficking to the Golgi apparatus. TMEM165, along with genes associated with heparan sulfate biosynthesis (SLC35B2, HS6ST1, NDST1), and Golgi trafficking (multiple COG complex subunits) were enriched in negative selected cells for both of our screens. These findings are consistent with prior genome-wide haploid screens that identified the same pathways as enriched in RSV protected populations, suggesting dependence on these pathways in multiple cell culture models [17]. Notably, Rab4a, a member of RAS GTPase superfamily known to regulate membrane trafficking, was a hit in the early infection screen, but not the viability screen. Consistent with our finding, an siRNA screen of nine Rab proteins revealed a statistically significant decrease in RSV viral RNA upon knockdown of Rab4a [42].

The pursuit of broad antiviral therapeutics largely relies upon the idea that there are common host factors or pathways that multiple viral families require over the course of infection. To contextualize our screening results, and identify commonalities across multiple viral families, we compared our findings with 29 published CRISPR screening datasets for viral infection. While several clusters of hits were shared amongst subsets of viruses, the RSV screens displayed relatively few overlaps. Those hits that did overlap belonged to COG complex members, golgi-related factors, and heparan sulfate biosynthesis. Beyond RSV, the collective analysis of these 29 screens suggests that discovery of truly ubiquitous host determinants for the purpose of developing pan-viral therapeutics will continue to be highly challenging. Regardless, the exercise of harmonizing multiple screens published by numerous other groups is a useful endeavor and the comparison of screening results across groups should be a standard practice for these investigations.

Overall, our work highlights the value of using both single-cell transcriptomics and functional genomics to comprehensively investigate RSV infection. During the critical first hours of infection, we produced a high-dimensional map of transcriptionally regulated pathways during RSV infection, thus providing insight into the antiviral response in infected and bystander cells. Furthermore, we utilized functional genomics to identify host factors important for RSV infection and contextualized our results with 29 other published CRISPR screens. Our data-rich study complements the existing RSV literature and will serve as a resource for ongoing investigations of this prolific pathogen.

## Limitations of Study

All experiments in this study were performed in A549 cells, a cell line derived from human lung. While A549 cells have intact interferon signaling, and are derived from respiratory cells, this cell line does not contain the structure, morphology or heterogeneous mixture of primary tissue. It will be valuable to further investigate the innate immune response to RSV infection and required host dependency factors in primary respiratory cells. Furthermore, our experiments were performed with one strain (A2) of RSV. Moving forward, it would be interesting to investigate host transcriptional similarities and differences after infection with different viral strains.

## Materials and Methods

### Cell Culture

HEp-2 (ATCC CCL-23) and A549 (ATCC CCL-185) cells were ordered from ATCC and cultured in DMEM supplemented with 10% FCS, penicillin, streptomycin, and glutamine. Cells were maintained at 37°C and 5% CO_2_. Cells were mycoplasma negative using the MycoAlert Mycoplasma detection kit (Lonza).

### RSV Propagation

For single-cell sequencing experiments, we used a genetically unmodified virus: Human respiratory syncytial virus (ATCC® VR-1540). For CRISPR screening experiments, we used Respiratory Syncytial Virus with EGFP (RSV-GFP5, ViraTree #R125). Both isolates of RSV were amplified on HEp-2 cells, PEG precipitated, pelleted through a sucrose cushion during ultracentrifugation, resuspended and stored at -80°C. Viruses were titered on HEp-2 cells using the standard TCID50 assay for ATCC® VR-1540 or viral focus assay for RSV-GFP5. Both viral isolates were mycoplasma negative using the MycoAlert Mycoplasma detection kit (Lonza).

### Single-Cell RNA sequencing

A549 cells were infected with RSV at an MOI of 0.3 in serum-free media. After a two hour incubation, cells were washed with serum-free media, and fresh media with serum was replenished. To reduce batch effects, cells were sequentially infected and harvested at the same time point for single-cell sequencing. Cells were trypsinized, murine NIH/3T3 (ATCC CRL-1658) were spiked in, and MULTI-seq sample barcoding was performed following the McGinnis et al. protocol [55]. Cells were subsequently counted and the manufacturer’s protocol for 10x Genomics 3’ v2 sequencing were performed. The protocol described in McGinnis et al. was followed for MULTI-seq library preparation. Samples were subsequently sequenced on the Illumina NovaSeq and aligned using 10x Genomics CellRanger v3.1.0 to hg38 and mm10 reference genome concatenated with the RSV genome. Sample barcodes were demultiplexed following previously published protocols (McGinnis et al.). Downstream analyses were performed using Scanpy 1.9.5 [56]. We filtered out cells that were two standard deviations below the mean total counts and mean total genes detected. To identify RSV infected cells, we quantified the viral reads present in our spike-in murine (3T3) control population. Infected cells were identified as cells with >40 viral UMIs, and uninfected bystander cells identified as cells with <30 UMIs. Differential expression was performed using MAST [20]. In order to assess interferon stimulation in infected and bystander cells, we scored single cells for ISG expression using the sc.tl.score_gene function from scanpy.

### Genome-wide CRISPR Screen

A549 cells were lentivirally transduced with Cas9-BLAST (Addgene 52962; gift from Feng Zhang) and blasticidin selected. These A549-Cas9 lines were subsequently transduced with the human GeCKO v2 library (Addgene 1000000049; gift from Feng Zhang) and puromycin selected. This library was subsequently infected with RSV-EGFP. EGFP negative or low cells were sorted on the Sony SH800 cell sorter. For the first iteration of the screen with a viability selection, cells were cultured for 1 week and surviving cells were collected for downstream analysis. For the second screen, EGFP negative and low cells were collected from the sorter separately. We subsequently extracted genomic DNA, amplified guide sequences, performed library prep and sequenced on the Illumina NextSeq as described in Wang et al. [29]. We performed enrichment analysis using the MAGeCK algorithm [23]. For the “early infection screen”, sequencing data from EGFP negative and low sorted groups were pooled for analysis.

### CRISPR Screen Comparative Analysis

Fastq files from all the published CRISPR screen datasets were processed using a standardized workflow centered around the MAGeCK program [PMID: 25476604]. The computer code for the workflow is available at https://github.com/czbiohub/CRISPRflow. For studies where fastq files were not available, MAGeCK gene summary result files were downloaded. Virus host factor genes were identified using a positive-selection FDR value cutoff of 0.01 in at least one of the CRISPR screens (29 published screens and two screens in this study). For Matia et al. (Vaccinia CRISPR screen), the lowest FDR among 27 screens varying virus mutant, MOI and number of reinfections were used in the hit selection process.

The results of our comparative screening analysis were visualized using a heatmap of - log10(FDR) values where rows represent genes and columns represent CRISPR screen datasets. Both the rows and columns were arranged by the hierarchical clustering algorithm using the “average” linkage method with the “cosine” distance metric.

The fastq files for the following CRISPR screen datasets were downloaded using links provided by the corresponding research articles: Diep et al. EV and RV [PMID:31527793], Han et al. H5N1(VN04_Low_) [PMID:29642015], Hoffmann et al. YFV and ZIKV [PMID:33338421], Liu et al. LCMV [PMID: 35502904], Marceau et al. DENV and HCV [PMID:27383987], Ooi et al. DENV-1-276RK, DENV-2-429557, DENV-3-Philippines/H871856 and DENV-4-BC287/97 [PMID:31384002], Wang et al. 229E, OC43 and SARS-CoV-2 [PMID:33333024], Flint et al. EBOV [PMID: 30655525], Baggen et al. 229E and SARS-CoV-2 (Low-stringency) [PMID 33686287], Daniloski et al. SARS-CoV-2 (MOI=0.01) [PMID: 33147445], Hein et al. HCMV (CRISPRn_screen_host) [PMID: 34697476], Schneider et al. HCoV-229E 33°C, HCoV-NL63 33°C, HCoV-OC43 33°C, SARS-CoV-2 33°C and SARS-CoV-2 37°C [PMID: 33382968], Qiao et al. PeV-A1 and PeV-A2 [PMID: 38658526], Richards et al. ReoT3D [PMID: 36074821].

## Acknowledgments

We thank Melanie Ott, Katrina Kalantar, Madhura Raghavan and the DeRisi lab for helpful discussions; Norma Neff and the CZ Biohub San Francisco Genomics Platform for sequencing support.

This work was supported by the Chan Zuckerberg Biohub San Francisco (J.L.D., A.S.P.) and the National Institute of Health (F31AI150007 to S.S.). C.J.Y. is supported by an NIAID R01 (R01AI171184), NIDCR U01 (U01DE028891), NIAID P01 (P01AI172523), and NHGRI UM1 (UM1HG012076). Additional support for C.J.Y. comes from NIAMS (P30AR070155), the Department of Defense, the Arc Research Institute, Parker Institute for Cancer Immunotherapy (PICI), and the Chan Zuckerberg Initiative. This work does not necessarily represent the official views of the NIH.

J.L.D. is a paid scientific advisor for Allen & Co. J.L.D. is a paid scientific advisor for the Public Health Company, Inc. and holds stock options. C.J.Y. is the founder of and holds equity in DropPrint Genomics (now ImmunAI) and Survey Genomics. C.J.Y. serves as a Scientific Advisory Board member for and holds equity in Related Sciences and ImmunAI, and is a consultant or has consulted for Maze Therapeutics, TReX Bio, HiBio, ImYoo, Kiragen, and Santa Ana Bio. Additionally, C.J.Y. is an Innovation Investigator at the Arc Institute. C.J.Y. has received research support from the Parker Institute for Cancer Immunotherapy, Chan Zuckerberg Initiative, Chan Zuckerberg Biohub, Genentech, BioLegend, ScaleBio, and Illumina.

## Data Availability

Raw and preprocessed single-cell data will be made available on GEO (GSE281623).

---

**Figure S1:**
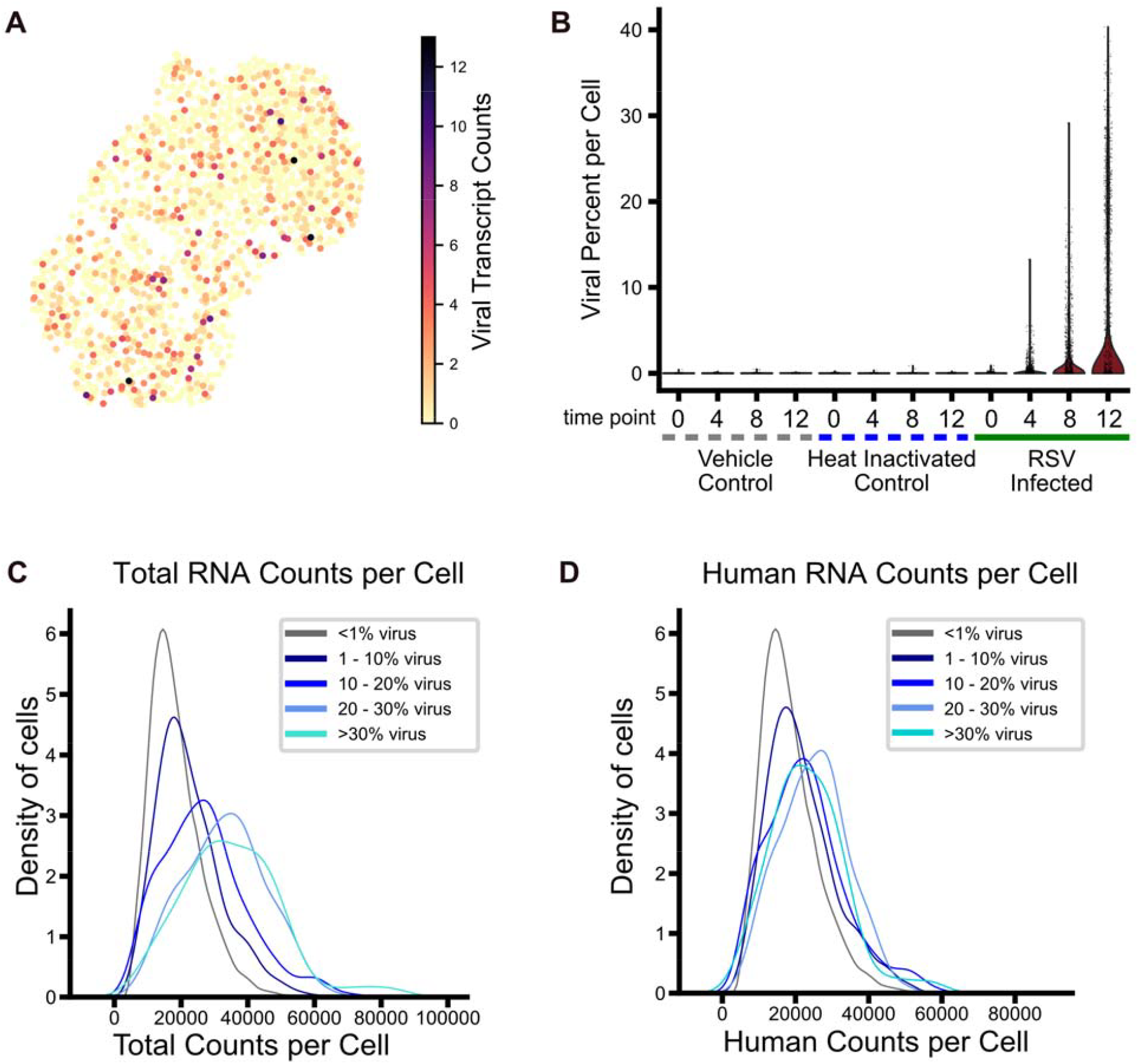
Evaluation of viral gene expression and distribution of RNA counts per cell. **A**. To evaluate ambient viral RNA captured in droplets during single-cell RNA sequencing, we included a spike-in of murine cells and quantified viral transcript counts per cell. **B**. The viral percentage per cell for each condition and time point was calculated. **C-D**. Cells were binned by percentage of viral transcripts and each plot represents the density for (**C**) total counts and (**D**) human counts per cell.

**Figure S2:**
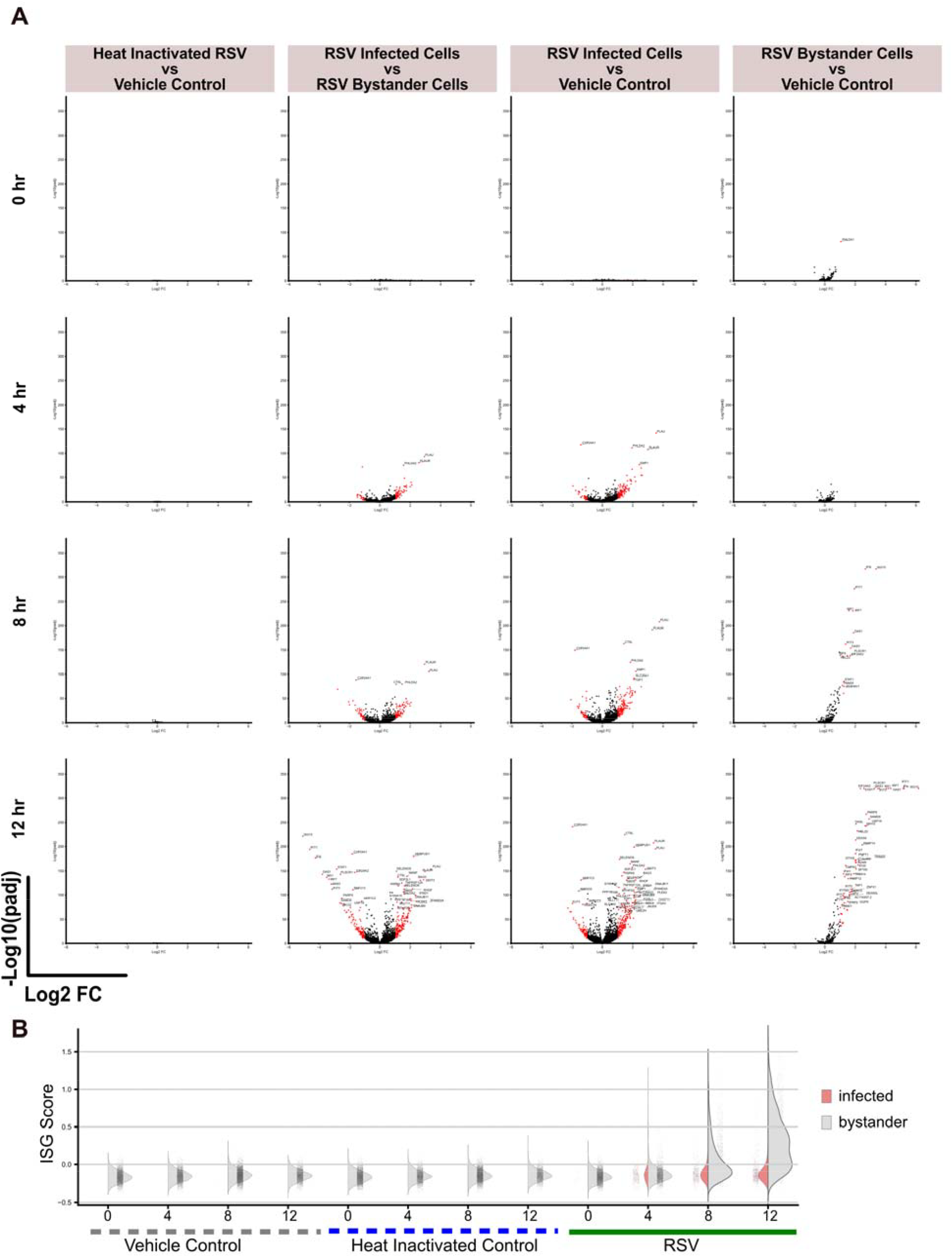
Differential expression results for each time point and condition. **A**. For each time point and treatment condition, we performed differential expression analysis (MAST). Resulting volcano plots show fold change (FC) and adjusted *P*-values for each pairwise comparison. **B**. Each cell was scored by expression of interferon-stimulated genes (ISGs), and ISG score for each time point and condition is shown for infected cells (pink) and bystander cells (gray).

**Figure S3:**
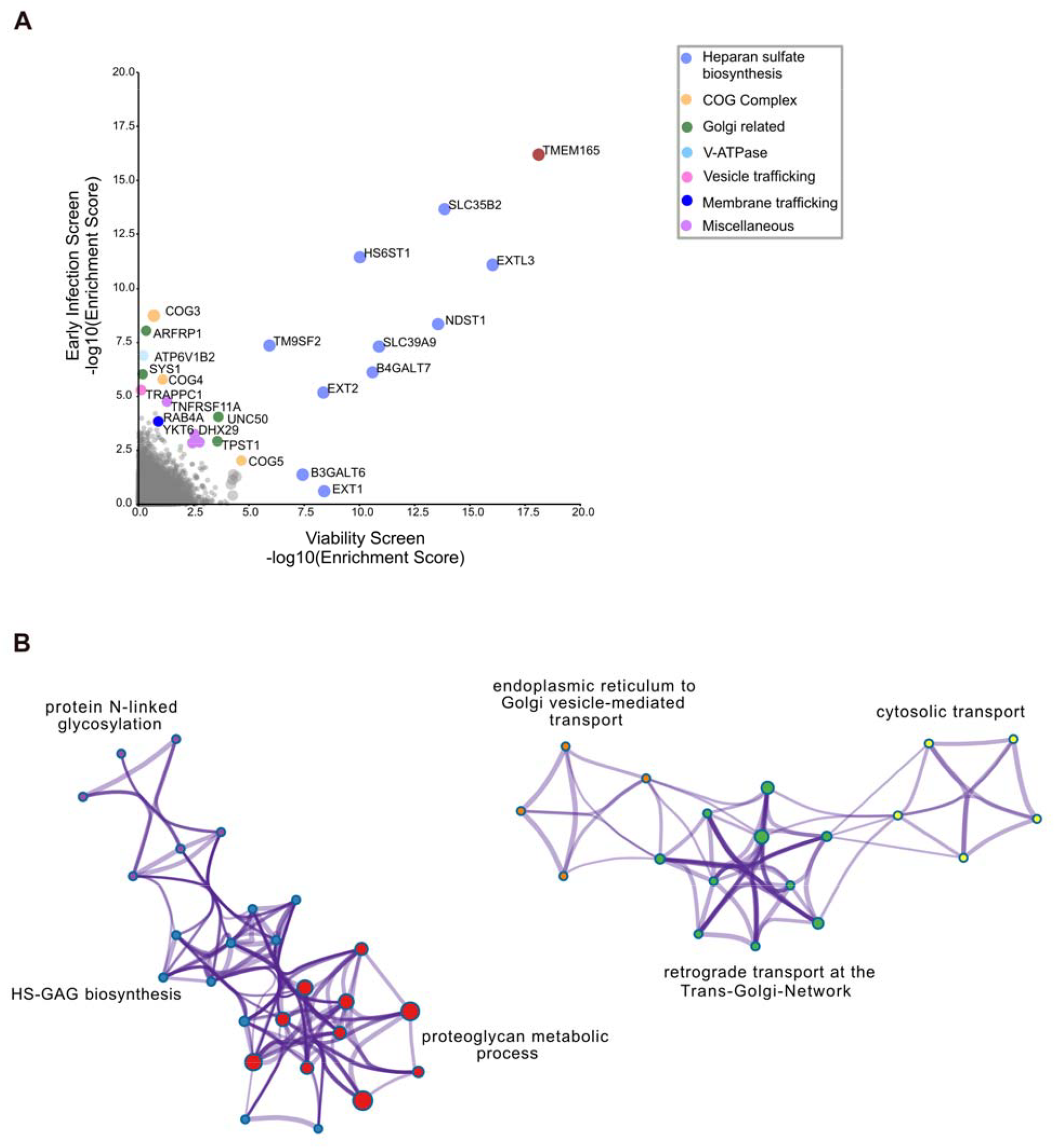
RSV screen comparisons and pathway enrichment of results. **A**. We compared the results of our two genome-wide screens using the -log10 MAGeCK enrichment scores. Each gene is a dot and top genes are colored by function. **B**. Pathway and process enrichment clusters from our top genes were identified using Metascape [27]. Each node in this network is an enriched term and nodes are colored by biological cluster description.

**Figure S4:**
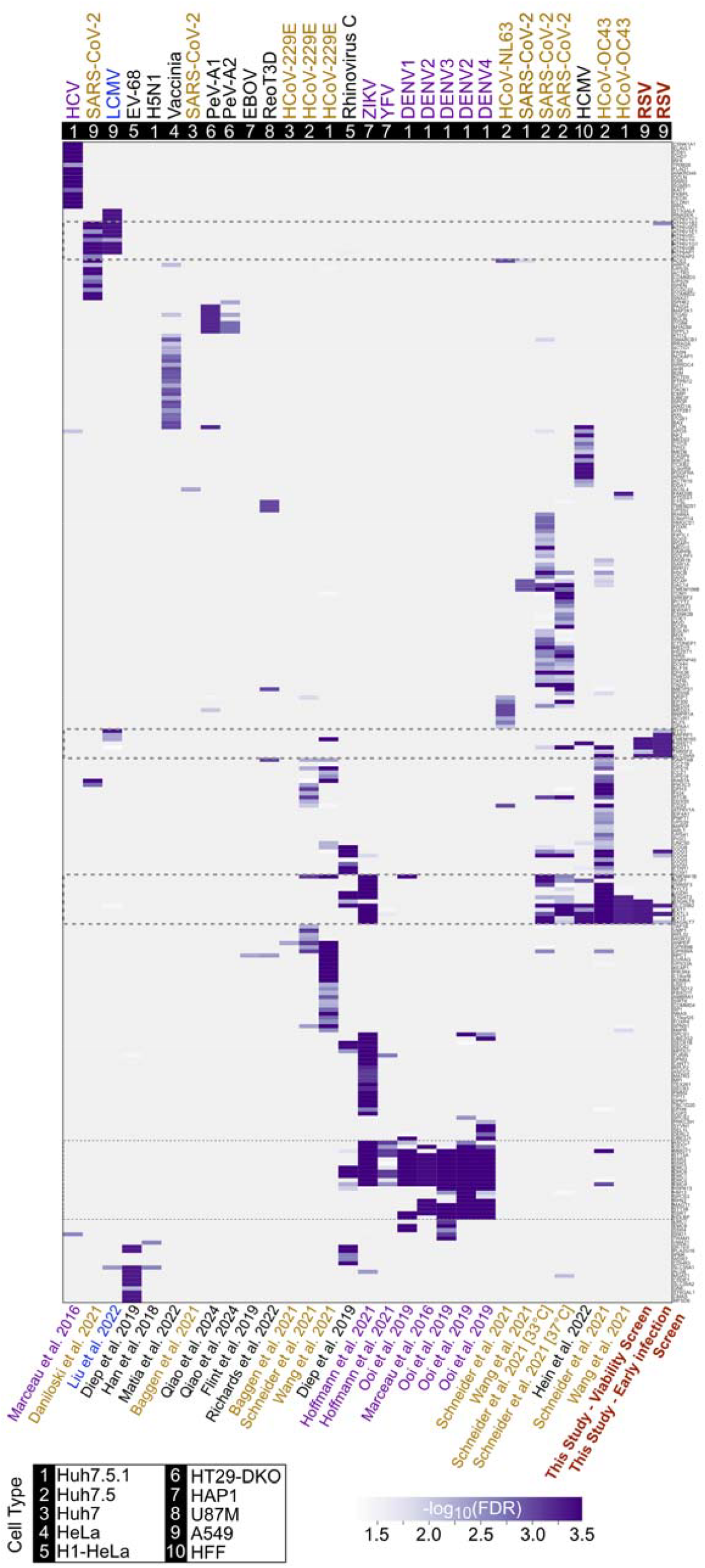
Contextualization of RSV screening results. Heatmap from Figure 4 detailed with all genes and associated study identifiers.

